# Strategy in wheat-*Fusarium* dual-genome RNA-seq data processing

**DOI:** 10.1101/2019.12.16.878124

**Authors:** Ziying Liu, Yifeng Li, Youlian Pan, Lipu Wang, Therese Ouellet, Pierre Fobert

## Abstract

In RNA-seq data processing, short read alignments are usually of one species against its own genome; however, in plant-microbe interaction systems, reads from both host and pathogen samples are blended together. In contrast to single-genome, both pathogen and host reference genomes are involved in the alignment process. In such circumstances, the order of alignment to the host, the pathogen, or simultaneously to both genomes results in differences in read counts of certain genes, especially at the advanced infection stage. It is crucial to have an appropriate strategy for aligning the reads to their respective genomes, yet the existing strategies of either sequential or parallel alignment become a problem when mapping mixed reads to their corresponding reference genomes. The challenge lies in the determination of which reads belong to which species, especially when homology exists between the two genomes.

This study proposes a combo-genome alignment strategy after comparing three alignment scenarios. Simulation results demonstrated that the degree of discrepancy in the results is correlated with phylogenetic distance of the two species in the mixture which was attributable to the extent of homology between the two genomes involved. This correlation was also found in the analysis using two real RNA-seq datasets of *Fusarium*-challenged wheat plants. Comparisons of the three RNA-seq processing strategies on three simulation datasets and two real *Fusarium*-infected wheat datasets showed that an alignment to a combo-genome, consisting of both host and pathogen genomes, improves mapping quality as compared to sequential alignment procedures.

## Introduction

High throughput RNA sequencing technology has become progressively accessible for functional genomics research and carries increased sensitivity and specificity compared to microarray technology (Nagalakshmi at al. 2008). A crucial task performed in RNA-seq data processing is aligning millions of short cDNA fragment (reads) to a reference genome. The percentage of uniquely mapped reads, a global indicator of the overall sequencing accuracy, is an important mapping quality indicator (Conesa et al. 2016). Since alignment results will affect subsequent data analysis, it is important to select an unbiased sequence alignment procedure.

In the RNA-seq processing procedure for a single species, RNA-seq reads are aligned to their reference genome directly. In a study on plant-microbial interaction, however, two genomes are involved; RNA-seq reads are thus a mixture from both pathogen and host genomes. Accurately determining whether the reads align to the host, the pathogen, or both genomes, is challenging when separately performing alignment to a single genome at a time.

Researchers use RNA-seq to study and monitor transcriptomic dynamics of host-pathogen interaction simultaneously under given conditions, and furthermore examine the strategies applied by the host and pathogen to compete for limited biochemical resources (Damron et al. 2016; Darren et al. 2017; Westermann et al. 2017). For example, Nuss and colleagues (2017) developed an experimental approach that allows simultaneous monitoring of genome-wide infection-linked transcriptional alterations of the host and pathogen. They first split the libraries into reads originating from mouse and pathogen (*Yersinia pseudotuberculosis* IP32953) using Bowtie and Tophat2, and then excluded the identified cross-mapped reads from downstream analysis. Finally, the split RNA-seq reads were aligned to their corresponding genome respectively. Westermann’s lab (2012) applied dual-genome RNA-seq data of *Salmonella*-infected human cells to monitor the host response to bacterial infection. The mixture of RNA-seq reads originated from either *Salmonella enterica* or hosts (HeLa cells from human or mouse) was mapped to their respective genomes with high stringency. Mapped reads, which could be aligned equally well to both host and *Salmonella* reference sequences, were discarded in subsequent analysis. Bradford and his colleagues (2013) developed a species-specific mapping workflow for RNA-seq data from xenograft. The RNA-seq data were aligned to the human and mouse genomes separately. To differentiate human and mouse expression, the reads that mapped to both human and mouse genomes were removed, which resulted in loss of yields in each species (2-2.5% for human and 17-26% for mouse). Damron et al. (2016) used dual RNA-seq to simultaneously measure *Pseudomonas aeruginosa* and the murine host’s gene expression in response to respiratory infection. Reads data were mapped to *P. aeruginosa* and *Mus musculus* genomes independently by using CLC Genomics platform. Dual genome RNA-seq approaches were also applied to monitor host-viral interactions (Woodhouse et al. 2010; Strong at al. 2013; Perez-Losada et al. 2015; Wesolowska-Andersen et al. 2017), assisting greatly in deciphering of viral-driven disease mechanisms.

In recent investigations into the transcriptomics associated with resistance and susceptibility of wheat against fusarium head blight (FHB), the RNA-seq data contain reads from both wheat and the pathogen, *Fusarium graminearum* (*F. graminearum*) (Pan et al., 2018, Wang et al., 2018, Fauteux et al., 2019). In this paper, we propose a combo-genome alignment approach which is to map RNA-seq data to a combination of both host and pathogen genomes. This approach was compared with two scenarios processing RNA-seq read alignment sequentially using three simulated datasets. Such comparison was also performed on two real-life datasets of *Fusarium*-infected wheat.

## Materials and Methods

### Simulated datasets

We selected *Arabidopsis thaliana* and three other species, with differences in their phylogenetic distances consisting of two dicots, *Arabidopsis lyrata* in the same genus, *Brassica rapa* in the same family, and one monocot *Brachypodium distachyon* (Fig. 1a). These four genomes were obtained from EnsemblPlants (http://plants.ensembl.org/, Release 33). *A. thaliana* was combined with each of the other three genomes, respectively. Each of the three combinations was used to randomly generate a 25 million paired-end RNA-seq dataset with 1% error rate, 101 bps in length by using the RNASeqReadSimulator tool (Li et al. 2012) (Fig. 1b).

**Fig. 1.**
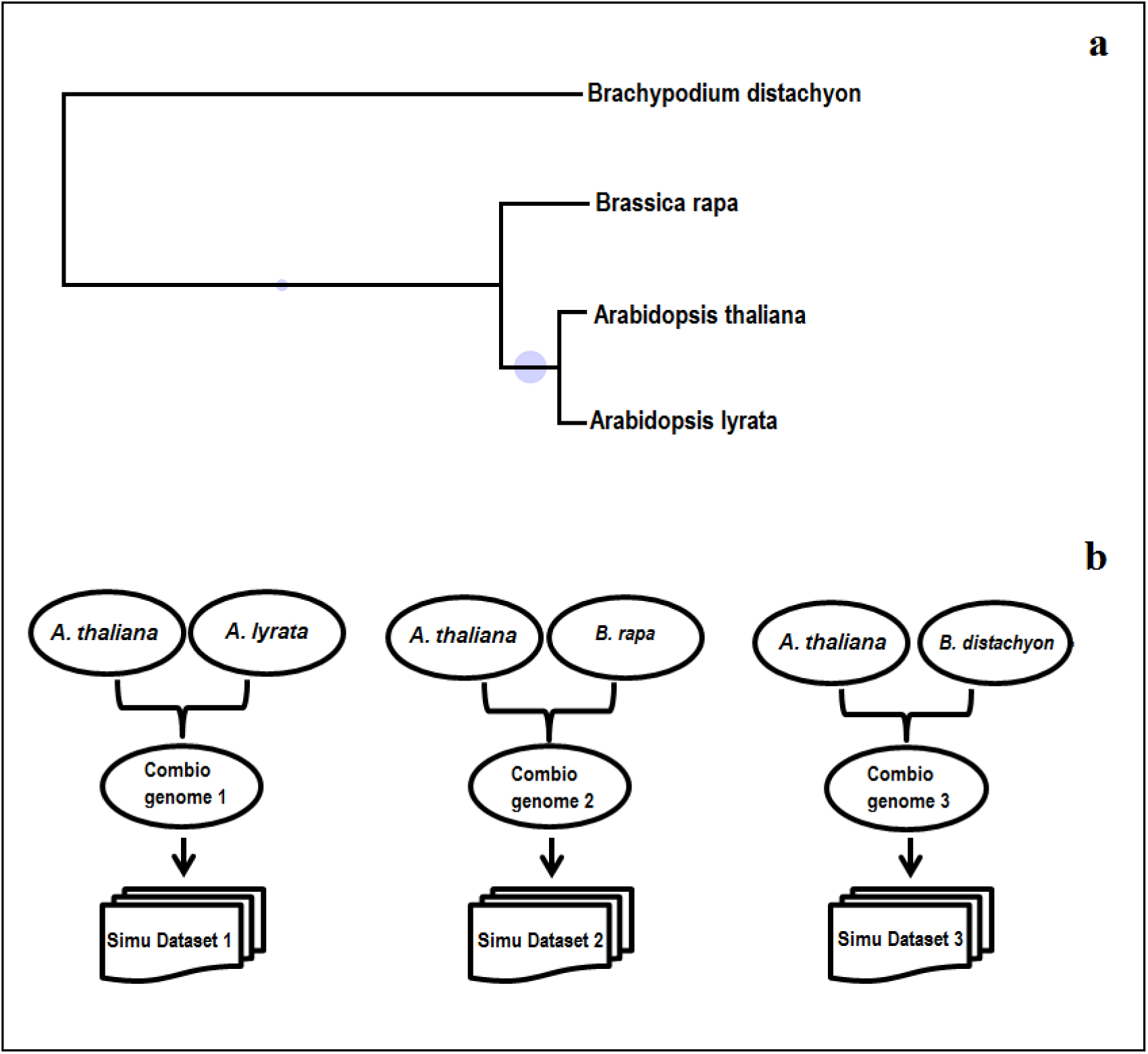
a) Phylogenetic distances between *A. thaliana* and each of three other species used for RNA-seq data simulation. Phylogenetic tree was created with iTOL (Letunic et al. 2011, https://itol.embl.de/). b) Our simulation process to generate dual-genome RNA-seq datasets.

### Real datasets

In addition to the simulated datasets, we processed two real datasets from FHB-infected wheat samples. Both datasets contain mixture of RNA-seq reads from wheat and *F. graminearum*. Each dataset includes a different set of four wheat genotypes of various resistance levels to FHB. There were three biological replicates for each sample. The first dataset consists of two resistant genotypes (Sumai3 and FL62R1) and two susceptible ones (Stettler and Muchmore); the samples were collected at 0, 1, 2 and 3 days post inoculation (dpi) (Wang et al. 2018, GSE118126). The second dataset consists of three resistant genotypes (Wuhan1, Nyubai and HC374) and one susceptible (Shaw) (Pan et al. 2018, GSE113128); the samples were collected at 2 and 4 dpi. We present the result of samples collected at 3 dpi (at longest dpi) in dataset 1 and 2 and 4 dpi in dataset 2 to illustrate our mapping strategy. IWGSC version 2.2 genome assembly (ftp://ftpmips.helmholtz-muenchen.de/plants/wheat/IWGSC/genome_assembly/genome_arm_assemblies_CLEANED_REPMASKED/ 2014) and MIPS version 2.2 IWGSC gene annotation (https://urgi.versailles.inra.fr/download/iwgsc/Gene_models/survey_sequence_gene_models_MIPS_v2.2_Jul2014.zip, 2014), fusarium (ftp://ftp.ensemblgenomes.org/pub/fungi/release-33/, 2016) were used in this study.

### Mapping Strategies

In dual-genome system, such as the host-pathogen system, two reference genomes and corresponding annotations were combined into a combo-reference genome and a combo-annotation, respectively.

After preprocessing of the RNA-seq reads to remove adaptor, low-quality reads and short reads (length<20 bps) using software package FASTX-toolkit-0.0.13 (http://hannonlab.cshl.edu/fastx_toolkit/), STAR (Dobin et al. 2013) was used to perform the alignments in the following three mapping strategies:

A. Dual-genome alignment: RNA-seq reads were aligned to the combo-genome with combo-annotation (Fig. 2a),
B. Sequential alignment 1: RNA-seq reads were aligned first to genome 1, then the unmapped reads were aligned to genome 2 (Fig. 2b), and
C. Sequential alignment 2: RNA-seq reads were aligned in reverse order of B), first to genome 2, then the unmapped reads were aligned to genome 1.

**Fig. 2.**
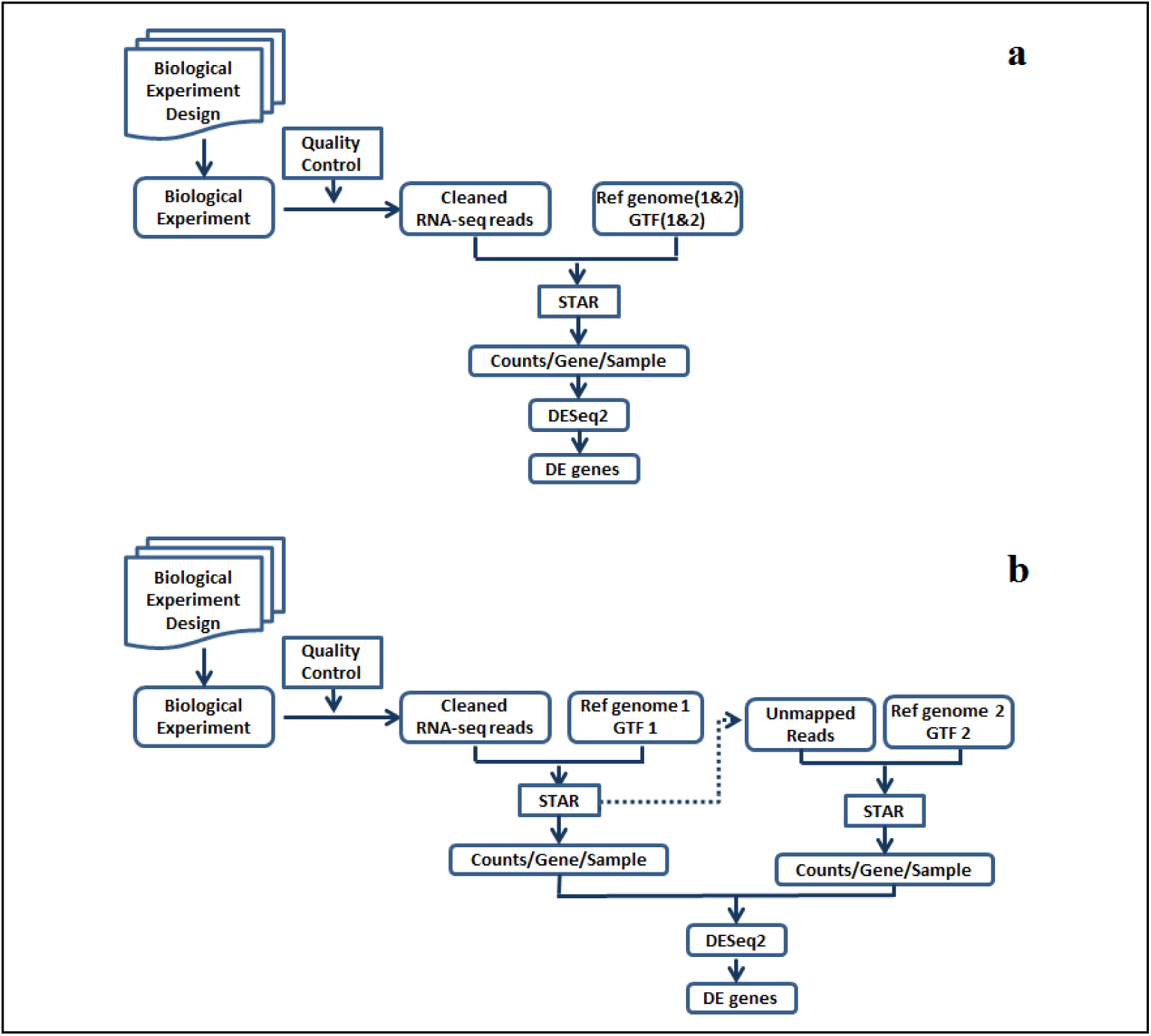
RNA-seq data analysis: a) Dual-genome alignment work flow, b) Sequential alignment work flow.

## Results

### Simulation results

Using the simulated datasets, our goal was to investigate the impact of homology to the mapping quality in dual-genome RNA-seq alignment. The mapping rate of sequential alignment varied significantly for the pairs of *A. thaliana* and *A. lyrata*, depending on the mapping order: 45.70% when *A. thaliana* is mapped first and 11.57% second, resulted in a difference of 34.13%. This difference decreased for more distant pairs between *A. thaliana* and *B. rapa* (4.28%). There was no visible difference disregarding the mapping order between the most distant pair *A. thaliana* and *B. discachyon* (Table 1).

**Table 1.**
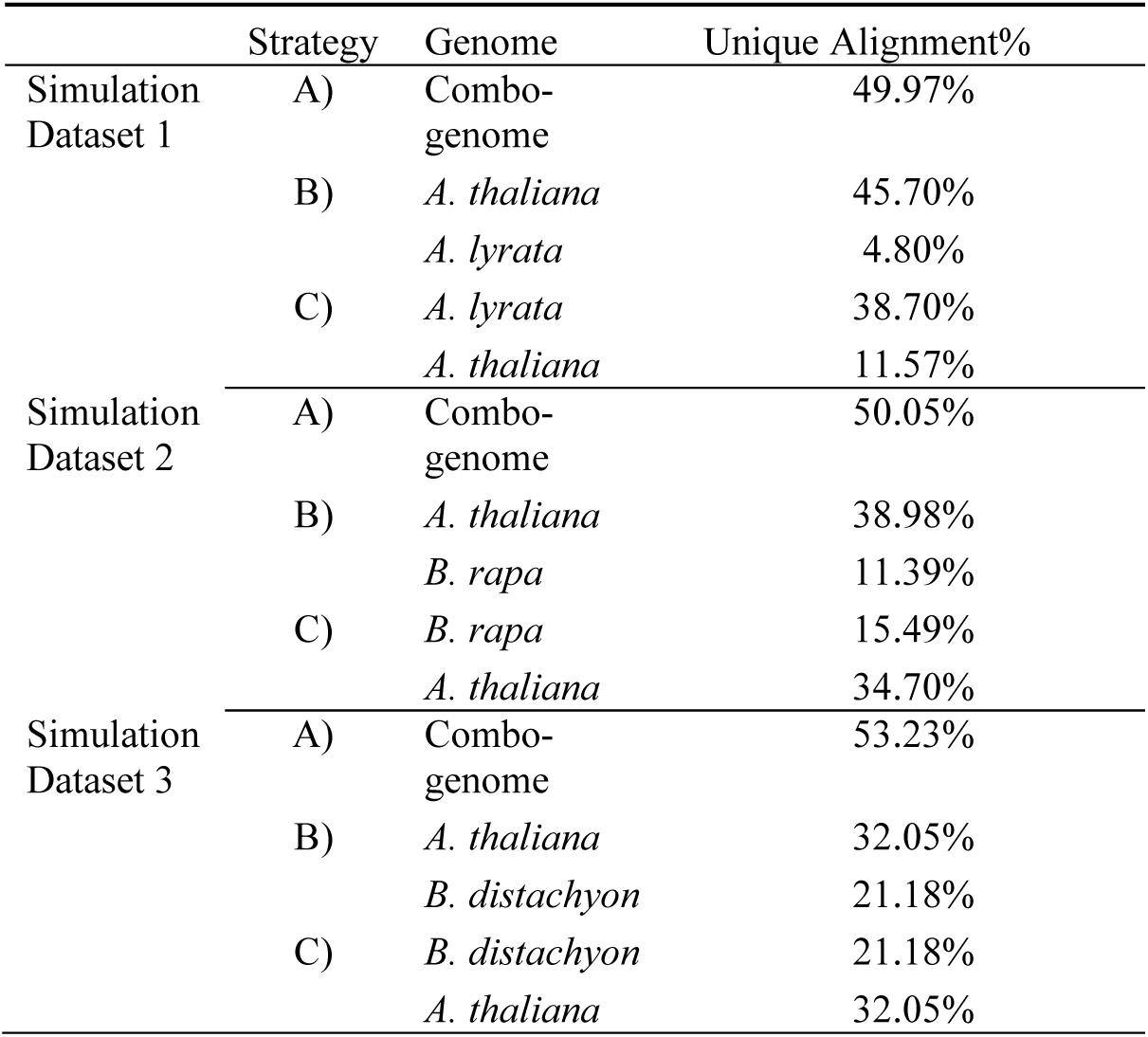
Simulation Results.

The difference in mapping rates between the two sequential alignments was correlated with the similarity of the genomes as represented by extent of homology between the two genomes (Fig. 3). The overall mapping rates of the combo-genome alignment were similar or equal to the sum of sequential alignments.

**Fig. 3.**
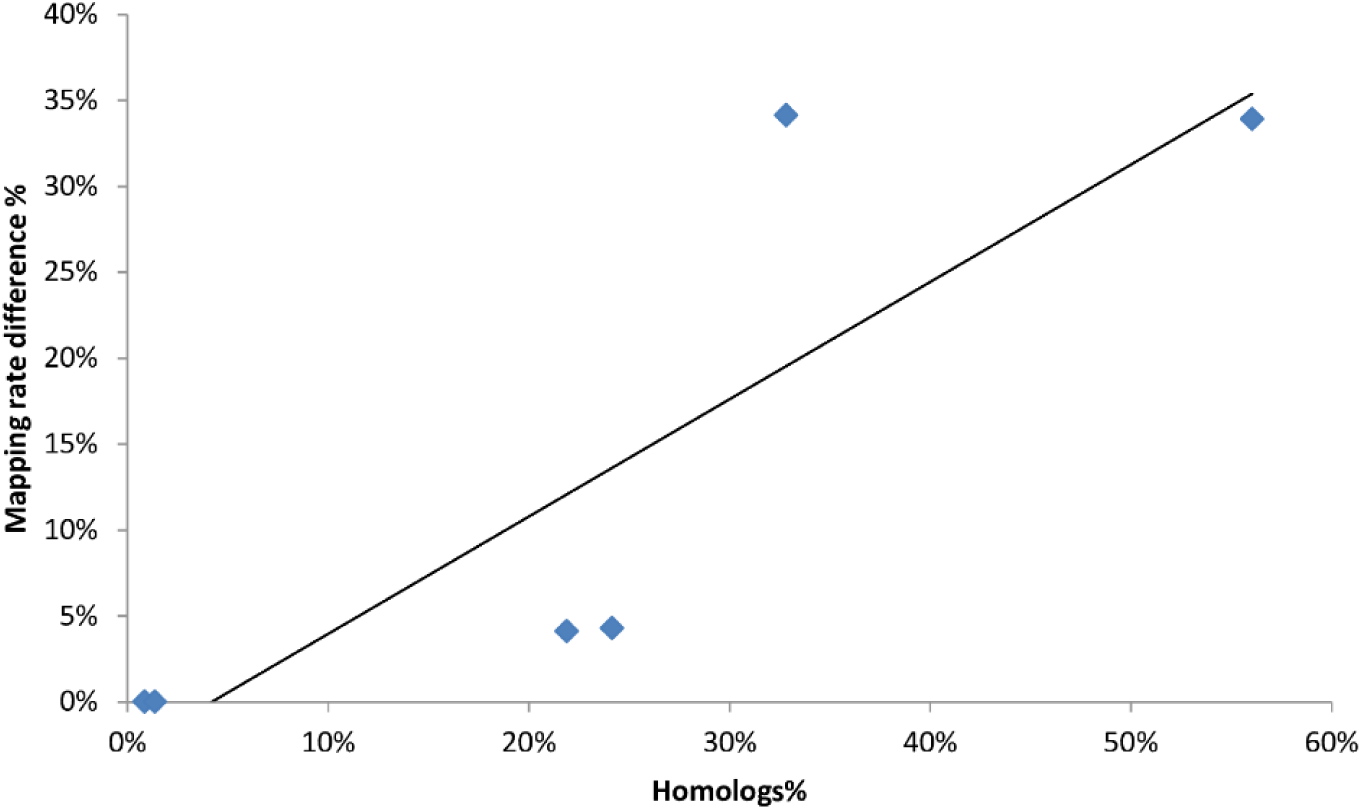
Correlation between percentage of homologs and the mapping rate difference between the two sequential strategies, r=0.85, p<0.05. Percentage of homologs between species was determined by reciprocal best hits using BLASTP.

### Real dataset results

On the two real datasets, the number of pathogen reads increases over time and was the highest at 3 (58.18%) and 4 dpi (65.19%) for Dataset 1 and Dataset 2, respectively (Fig. 4). The number of pathogen reads in resistance genotypes is much less than that in susceptible genotypes (Fig. 4b).

**Fig. 4.**
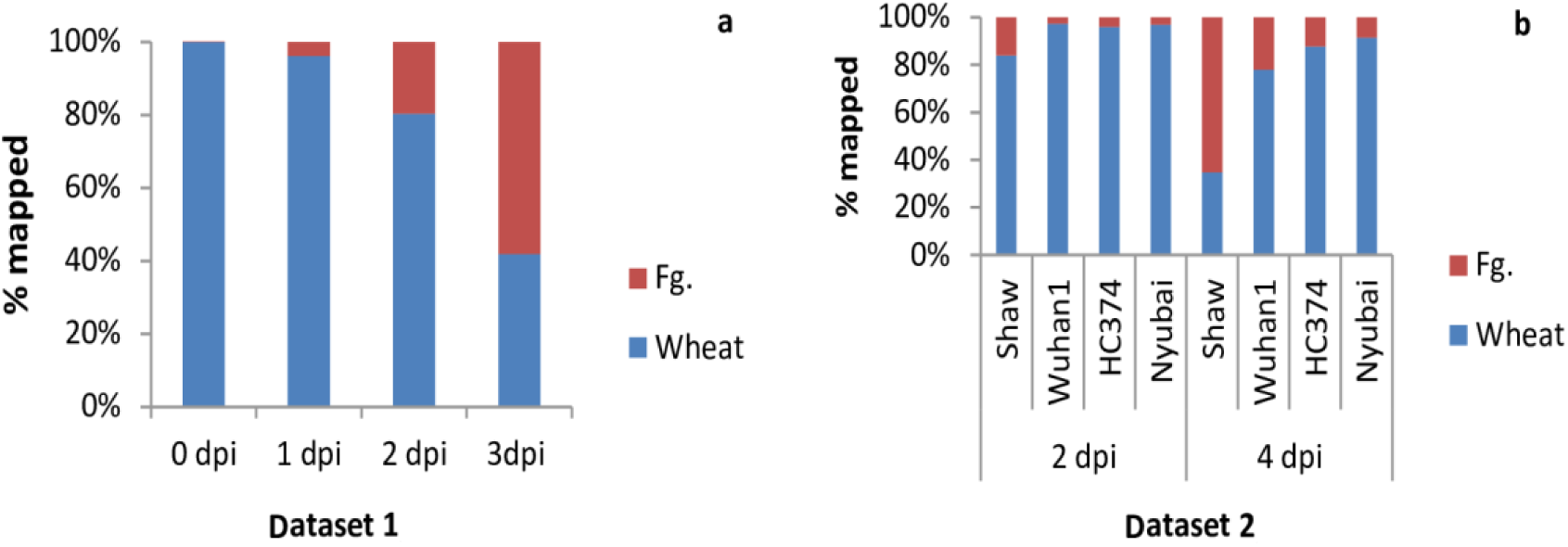
Dynamic interactome between wheat and *F. graminearum* (Fg) using results from strategy A. a) in the first three days after *F. graminearum* inoculation in resistance genotype Sumai3; b) from susceptible (Shaw) to resistance genotypes at 2 and 4 dpi.

The mapping results vary in the sequential alignment depending on the alignment order: host first (strategy B) or pathogen first (strategy C). The differences of the mapping rate are showed in Fig. 5.

**Fig. 5.**
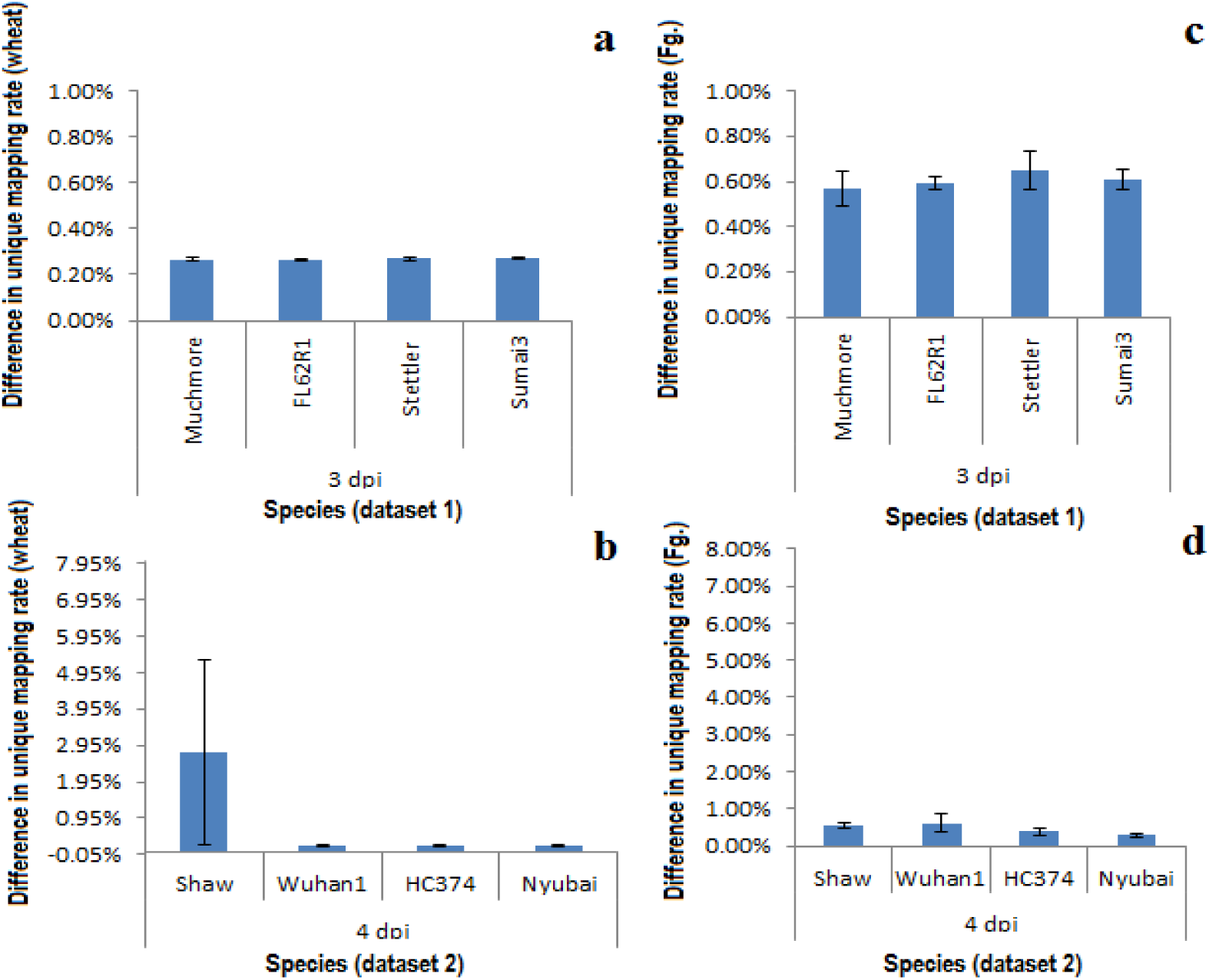
Differences in unique mapping rates between two sequential alignment strategies; **a**) and **b**) are the differences in wheat unique mapping between strategies B and C for datasets 1 and 2 respectively; while **c**) and **d**) are differences in *F. graminearum* unique mapping. Error bar indicates standard error of the mean.

The differences in unique mapping rate between strategies A (combo-genome strategy) and B (wheat first), and between strategies A and C (*F. graminearum* first) are shown in Fig. 6. The significant difference in unique mapping rates between strategies A and B is consistent on the two datasets.

**Fig. 6.**
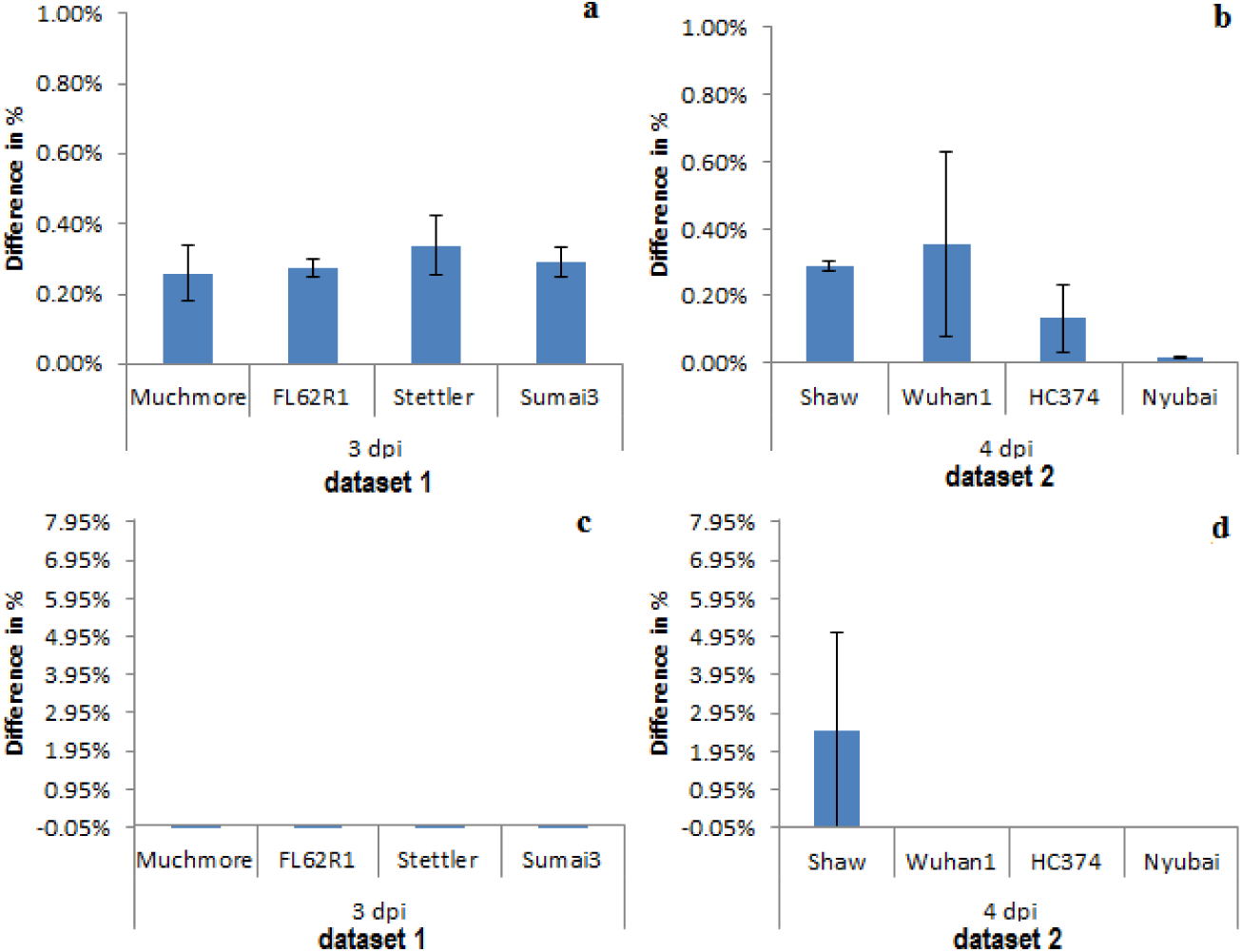
Differences in unique mapping rates between the combo-genome alignment strategy and two sequential alignment strategies; **a**) and **b**) are the differences between strategies A and B for datasets 1 and 2 respectively; while **c**) and **d**) are differences between strategies A and C for datasets 1 and 2 respectively. Error bar indicates standard error of the mean.

## Discussion

In this paper, we introduced a strategy of aligning the RNA-seq reads to the combo-genome references, combining both the host and the pathogen reference genomes, and compared its results with sequential alignment strategies. In the case of a pathogen-host system, the alignment of the reads to the factual genome is difficult because it is not practical to isolate the host RNA-seq reads from the pathogen completely, and homolog genes exist between the two genomes. In a sequential alignment, the mapping result is strongly biased toward the first reference genome in the processing order (Fig. 5). The extent of the bias depends on the proportion of homology between the two genomes, as demonstrated from the simulation results (Fig. 3): the higher the number of homologs between the two genomes is, the higher number of reads would be mapped to the first genome, leading to lower mapping rate to the second genome. Callari and colleagues (2018) developed a method using *in silico* combined human-mouse reference genome for alignment discriminates between human and mouse reads in patient-derived tumour xenograft to reduce this bias. The difference of unique mapping rate becomes insignificant when there are fewer homologs between the genomes such as in the pairs of *A.thaliana* and *B.distachyon*. However, the difference between strategies B and C for these real datasets is less significant than for the simulated datasets; this is due to the simulated datasets being generated using species with a much larger number of homologs for experimental purpose. This difference appears to be trivial for the real datasets; however, this bias will consequently translate to errors in the subsequent data quantification, normalization and differentially expressed gene identification. The intention of the combo-genome strategy introduced in this study is to mitigate such bias.

The alignment results for real datasets demonstrated significant improvement of unique mapping rates in strategy A as compared with strategy B. When compared with strategy C, we observed a mapping gain by using strategy A, in the mapping of susceptible genotype at 4 dpi when the wheat is severely infected by *F. graminearum* (Fig. 6). The differences on other genotypes were marginal; this is attributable to (1) the small *F. graminearum* genome as compared with the huge wheat genome, and (2) the number of actual *F. graminearum* reads is very small in the resistant genotypes. When mapping to the combo-genome, various possible alignments are considered and the optimal one is usually picked, i.e. a short read will be aligned to the most matched loci. For alignment of the similar quality reads, the mapping to both references equally well will be reported as multi-mapping reads (Dobin et al. 2013). This can help reduce the false positives and false negatives for unique calls to each species when compared with sequential alignment. In this way, less information will be lost. As such we would expect that the combo-genome strategy offers a more reliable alignment result. Our results for *F. graminearum* infected wheat datasets demonstrate that the combo-genome alignment strategy provided a higher quality mapping for dual-genome RNA-seq data. We believe that this strategy is universally applicable to other pathogen-host dual-genome alignment circumstances by mitigating the bias produced by separate mapping approaches due to the homology between the two genomes. We thus conclude that the combo-genome approach outperform sequential mapping strategy in dual-genome RNA-seq system.

## Acknowledgments

We are thankful to Linda Harris from Agriculture and Agri-Food Canada (AAFC) and Sean Walkowiak from Canadian Grain Commission for their insightful discussion at the early stage of this project.

